# Gill ionocyte remodeling mediates blood pH regulation in rockfish (*Sebastes diploproa*) exposed to environmentally relevant hypercapnia

**DOI:** 10.1101/2024.05.13.593928

**Authors:** Garfield T. Kwan, Alexander M. Clifford, Kaelan J. Prime, Till S. Harter, Martin Tresguerres

**Affiliations:** Marine Biology Research Division, Scripps Institution of Oceanography, University of California San Diego, USA; NOAA Fisheries Service, Southwest Fisheries Science Center, USA

**Keywords:** Ocean acidification, acidosis, upwelling, transcriptomics, slc9a3

## Abstract

Marine fishes excrete excess H^+^ using basolateral Na^+^/K^+^-ATPase (NKA) and apical Na^+^/H^+^-exchanger 3 (NHE3) in gill ionocytes. However, the mechanisms that regulate H^+^ excretion during exposure to environmentally relevant hypercapnia (ERH) remain poorly understood. Here, we explored transcriptomic, proteomic, and cellular responses in gills of juvenile splitnose rockfish (*Sebastes diploproa*) exposed to three days of ERH conditions (pH ∼7.5; ∼1,600 μatm *p*CO_2_). Blood pH was fully regulated at ∼7.75 despite a lack of significant changes in gill (1) mRNAs coding for proteins involved in blood acid-base regulation, (2) total NKA and NHE3 protein abundance, and (3) ionocyte density. However, ERH-exposed rockfish demonstrated increased NKA and NHE3 abundance on the ionocyte plasma membrane coupled with wider apical membranes and greater extension of apical microvilli. The observed gill ionocyte remodeling is consistent with enhanced H^+^ excretion that maintains blood pH homeostasis during exposure to ERH and does not necessitate changes at the expression or translation levels. These mechanisms of phenotypic plasticity may allow fishes to regulate blood pH during environmentally relevant acid-base challenges, and thus have important implications for both understanding how organisms respond to climate change and for selecting appropriate metrics to evaluate its impact on marine ecosystems.

## Introduction

Many marine environments regularly experience elevated *p*CO_2_ levels (“hypercapnia”) of variable magnitude and duration. For instance, *p*CO_2_ in kelp forests, coral reefs, and tide pools can easily reach up to 1,000 μatm (pH ∼7.70) on a nightly basis due to the respiration of residing organisms (reviewed in Tresguerres et al 2020). Upwelling events can induce surges in *p*CO_2_ that reach levels higher than 1,200 μatm (pH ∼7.6) at the ocean surface for several days to weeks (Chavez et al., 2018; Frieder et al., 2012; Hamilton et al., 2017; Hofmann et al., 2011; Kroeker et al., 2023; Takeshita et al., 2015). On a longer time scale, anthropogenic ocean acidification (OA) is predicted to increase global average surface ocean *p*CO_2_ levels to ∼1,000 μatm *p*CO_2_ by the year 2100, and to ∼2,000 μatm *p*CO_2_ (pH ∼7.4) by the year 2300 (Goodwin et al., 2018).

Exposure to hypercapnia results in a blood acidosis in aquatic animals. To avoid negative effects on gas transport and other physiological functions, marine fishes mitigate blood acidosis using specialized gill ion-transporting cells called “ionocytes”. These cells express abundant basolateral Na^+^/K^+^-ATPase (NKA) that generates the driving force for excreting H^+^ into seawater in exchange for Na^+^ through apical Na^+^/H^+^ exchangers 3 (NHE3; slc9a3) (Hiroi and McCormick, 2012). It is well established that marine fishes chronically exposed to severe hypercapnia (>6,000 μatm *p*CO_2_; pH <7.00) increase the abundance of gill NKA and NHE3 mRNA and protein (Deigweiher et al., 2008; Edwards et al., 2005; Melzner et al., 2009a), which results in greater H^+^ excretion from the blood into the seawater to restore nominal blood pH levels (Edwards et al., 2005; Liu et al., 2016).

In contrast, the mechanisms that upregulate H^+^ excretion during exposure to environmentally relevant hypercapnia (ERH) are less clear. Only a few studies have studied the gill ion-transporting mechanisms underlying blood pH regulation in marine fishes exposed to ERH. The Gulf toadfish (*Opsanus beta*) regulated blood pH during 3-day exposures to 1,900 μatm *p*CO_2_ (Esbaugh et al., 2012). Unlike the responses of marine fishes to chronic severe hypercapnia, toadfish gill NKA activity significantly decreased after 24h of exposure to ERH and then it reverted back to control levels by 72 h (Esbaugh et al., 2012). Atlantic cod (*Gadus morhua*) upregulated the mRNA abundance of three gill NKA α subunit isoforms after a 28-day exposure to 1,200 μatm *p*CO_2_ (but not to more severe levels of 2,200 μatm *p*CO_2_); however, gill NKA protein abundance and activity remained unchanged (but blood pH was not measured) (Michael et al., 2016). Red drum (*Sciaenops ocellatus*) fully regulated blood pH after a 14-day exposure to 1,000 μatm *p*CO_2_ and significantly increased gill NKA activity (Esbaugh et al., 2016) and a follow up study did not detect any changes in NHE3 mRNA abundance under comparable conditions (Allmon and Esbaugh, 2017). European seabass (*Dicentrarchus labrax*) also regulated blood pH during exposure to 1,000 μatm *p*CO_2_, where gill NKA mRNA expression remained unchanged over the first three and seven days, but upregulated after 21 days of hypercapnic exposure (though protein abundance was not measured) (Shrivastava et al., 2019). Altogether, these studies indicate that marine fishes can fully regulate blood pH during ERH exposure. However, the experimental differences between studies, the variable and small changes in gill NKA and NHE3 abundances, and the disconnect between mRNA expression and protein abundance currently do not allow clear generalizations on the physiological mechanisms of plasticity that regulate blood pH during ERH.

More recently, it was reported that European seabass exposed to 10,000 μatm *p*CO_2_ (pH ∼6.90) were able to regulate blood pH within 2 h (Montgomery et al., 2022). These fish did not experience any changes in gill NKA or NHE3 protein abundance; however, immunohistochemical observations revealed that the gill NKA-rich ioncytes had significantly increased apical surface area with NHE3 on it. These results indicated blood pH regulation was achieved *via* activation of pre-existing gill NKA and NHE3 proteins. However, it is not clear whether this mechanism is specific to European seabass, or whether it is relevant during longer exposures to EHR.

In addition to NKA and NHE3, the V-type H^+^-ATPase (VHA) has been proposed to excrete H^+^ to compensate a blood acidosis in marine fishes, as it is the case in freshwater fishes (Tresguerres et al., 2023). Although VHA mRNA abundance was upregulated in some cases during exposure to ERH, there was little correlation between VHA mRNA expression and protein abundance (Allmon and Esbaugh, 2017; Michael et al., 2016) and VHA was not observed in the apical membrane of gill cells (Allmon and Esbaugh, 2017), where it would be expected to be found should it play a role in H^+^ excretion.

We recently reported that juvenile splitnose rockfish (*Sebastes diploproa*) have two types of gill ATPase-rich cells. One is the archetypical NKA-rich ionocyte, which additionally expressed VHA throughout its cytoplasm. The other one has abundant VHA in its basolateral membrane, does not express detectable NKA, and its function is still unclear (Tresguerres et al., 2023). We also reported that these fish were able to fully regulate blood pH upon exposure to ERH (1,600 μatm *p*CO_2_, pH ∼7.5, for three days) and identified the physiological mechanism that links blood and endolymph acid-base regulation with otolith overgrowth (Kwan and Tresguerres, 2022). In the current paper, we examined the gills from those same rockfish to investigate the molecular and cellular mechanisms underlying blood acid-base regulation in response to ERH. Specifically, we analyzed the following in splitnose rockfish gill: 1) transcriptomic responses using RNAseq, 2) NKA, NHE3, and VHA protein abundance in crude homogenates and membrane-enriched fractions using Western blotting, 3) NKA-rich ionocyte and VHA-rich cell abundance, and NKA, NHE3, and VHA subcellular localization using immunohistochemistry and confocal microscopy, and 4) NKA-rich ionocyte apical morphology using scanning electron microscopy (SEM). Our integrative physiological genomics approach indicated that the full compensation of rockfish blood pH during ERH involves a functional remodeling of gill ionocytes by translocating existing NKA and NHE3 proteins into the plasma membrane of ionocytes together with an extension of apical microvilli that collectively upregulated H^+^ excretion into seawater.

## Methods

### Experimental animals, tissues, and setup

The gill samples analyzed in this study were dissected from the same splitnose rockfish reported in Kwan and Tresguerres (2022). Rockfish were caught from drifting kelp paddies off the coast of La Jolla and raised in the Hubbs Experimental Aquarium at Scripps Institution of Oceanography at the University of California San Diego (SIO-UCSD, La Jolla, USA; California Department of Fish and Wildlife permit #SCP13227). Rockfish average length and weight were 12.08 ± 0.18 cm and 45.54 ± 1.89 g (mean ± SEM; N=21), respectively.

Briefly, two header tanks were supplied with ambient seawater: the control header tank was not manipulated and its *p*CO_2_ and pH were 571.90 ± 4.88 µatm and 7.89 ± 0.012, respectively, whereas the ERH header tank was bubbled with CO_2_ using a pH-stat system (IKS Aquastar, Karlsbad, Germany) to adjust *p*CO_2_ and pH to 1,591.56 ± 18.58 µatm and 7.49 ± 0.01, respectively. Header tank supplied water flowed into 3-L experimental tanks (flow rate = 0.3 L/min) where rockfish were individually housed. After a 12 h acclimation period, fishes were exposed to control or ERH treatment conditions for 72 h. Rockfish were not fed throughout the experiment to ensure similar metabolic states. Experiment was repeated three times, with three controls and three ERH-exposed rockfish. No mortality was observed.

Blood and gill sampling, blood pH and plasma total CO_2_ measurements, and plasma *p*CO_2_, [HCO_3_ ^-^] and [CO_3_^2-^] calculations were performed as described in Kwan and Tresguerres (2022). A blood aliquot was loaded in a capillary tube, centrifuged (500 g, 3 min) and used to determine Hct. After euthanasia, gill samples were dissected and fixed for immunohistochemistry (1^st^ right arch) or SEM (1^st^ left arch), immersed in RNAlater for RNAseq analysis (2^nd^ right arch), or flash frozen in liquid nitrogen for Western blot analysis (2^nd^ left arch). All experiments were approved by SIO-UCSD animal care committee under the Institutional Animal Care and Use Committee protocol (#S10320).

### RNA extraction, sequencing and analysis

Gill samples were incubated in RNAlater for 24 hours at 4°C, then stored at −20°C until processing. Frozen samples were then thawed on ice, blotted dry, transferred to a 2 mL tube containing TRIzol® Reagent, and homogenized on ice with a handheld motorized mortar and pestle (Kimble®/Kontes, Dusseldorf, Germany). The TRIzol homogenate was briefly centrifuged to pellet debris, and the supernatant was processed for RNA extraction (RNAEasy Mini; Qiagen) according to the manufacturer’s specifications with an on-column DNAase I treatment to digest trace gDNA. RNA quantity and purity were determined via spectrophotometry (Nanodrop; Thermo Scientific); RNA integrity was determined using an Agilent 2100 Bioanalyzer (Agilent, Santa Clara, CA, USA). High-quality RNA samples (RNA Integrity Number > 8) were used to construct individual Poly-A enriched cDNA libraries by Novogene (Beijing, China) using the Illumina TruSeq RNA Sample Preparation Kit (Illumina, San Diego, CA, USA) according to the manufacture’s protocol. Briefly, mRNA was purified from total RNA using Oligo(dt) magnetic beads and the retained mRNA was chemically sheared into shorter fragments, followed by cDNA synthesis using random hexamer primers. The cDNA was fitted with Illumina adaptor primers and subjected to end-repair processing. After agarose gel electrophoresis, 200-300 bp fragments were extracted and used as templates for PCR amplification and cDNA library preparation. Libraries were sequenced using Illumina NovoSeq™ 6000 platform to generate unstranded 150 bp paired-end reads (n = 6; Q30 > 83%). Raw sequencing reads were analyzed, trimmed of adaptor sequences, and processed to remove i) low quality reads (qualified quality phred score < 20), ii) reads containing >50% unqualified bases (Base quality < 5), and iii) reads possessing >10 unknown bases (Ns) with fastp (Chen et al., 2018). Remaining unpaired reads were discarded from downstream analysis. Standard RNAseq quality control metrics are shown in Table S1.

*S. diploproa* gill quality reads were aligned and mapped to the published genome of honeycomb rockfish *S. umbrosus*) (NCBI Accession: PRJNA562005; fSebUmb1.pri; downloaded November 3, 2020) using HISAT2 (V2.1.0; Kim et al., 2015). Mapped reads were assembled into transcripts, merged amongst samples, and this merged transcript file was used to quantify gene and transcript abundance in each sample using Stringtie (V2.1.2; Pertea et al., 2015; Kovaka et al., 2019). Final gene and transcript quantitation data were analyzed with *R* using DESeq2 (Love et al., 2014). Genes of low abundance (average abundance <1 across all samples) and very-low expression (summed counts < 5 across all samples) were filtered from the expression matrix. Following initial DESeq2, expression data was again normalized to remove differences across technical sources of variation using RUVseq (Risso et al., 2014) and data was processed again using DESeq2.

### Antibodies

NKA was immunodetected using a mouse monoclonal antibody against the α-subunit of chicken NKA (a5, Developmental Studies Hybridoma Bank, Iowa City, IA, USA; Lebovitz et al., 1989). NHE3 was immunodetected using custom-made rabbit antibodies against two epitope regions within rainbow trout (*Oncorhynchus mykiss*) NHE3b (GDEDFEFSEGDSASG and PSQRAQLRLPWTPSNLRRLAPL) (Christensen et al., 2012). VHA was immunodetected using custom-made rabbit polyclonal or mouse monoclonal antibodies (GenScript, Piscataway, USA) against a highly conserved epitope within the VHA subunit B (AREEVPGRRGFPGY) (Kwan et al., 2020; Yee et al., 2019). These three primary antibodies were validated in splitnose rockfish (Figure S1 and Kwan and Tresguerres 2022). The secondary antibodies were goat anti-mouse HRP-linked and goat anti-rabbit HRP-linked (Bio-Rad, Hercules, CA, USA) for immunoblotting, and goat anti-rabbit Alexa Fluor 488 and goat anti-mouse Alexa Fluor 568 (Invitrogen, Grand Island, USA) for immunohistochemistry.

### Western Blotting

Frozen gill samples were immersed in liquid nitrogen, pulverized using a porcelain mortar and pestle, and submerged within an ice-cold buffer containing protease inhibitors (250 mmol l^−1^ sucrose, 1 mmol l^−1^ EDTA, 30 mmol l^−1^ Tris, 10 mmol/L benzamidine hydrochloride hydrate, 1 mmol/L phenylmethanesulfonyl fluoride, 1 mmol/L dithiothreitol, pH 7.5). Samples were centrifuged at 3,000xg (10 min, 4°C) to remove cell debris, and the resulting supernatant was saved as the “crude homogenate”. A portion of the crude homogenate was further centrifuged at 21,130xg (30 min, 4°C), and the pellet was saved as the “membrane-enriched” fraction. Total protein concentration in both fractions was determined using the Bradford assay (Bradford, 1976), and stored at −80°C if not used on the same day. On the day of SDS-electrophoresis, the samples were mixed with an equal volume of 2x Laemmli buffer containing 10% β-mercaptoethanol, heated (70°C for 5 min) and loaded (crude homogenate: 10 µg per lane; membrane: 2 µg per lane) onto a 7.5% polyacrylamide mini gel (Bio-Rad, Hercules, CA, USA). Proteins in the samples were separated by polyacrylamide gel electrophoresis at 200 V for 40 min and the separated proteins were transferred onto a polyvinylidene difluoride (PVDF) membrane using a wet transfer cell (Bio-Rad) at 100 mAmps (4°C overnight). On the following day, PVDF membranes were incubated in tris-buffered saline with 1% tween (TBS-T) with skim milk powder (0.1 g/mL) for 1 h, then incubated with primary antibody (NKA: 10.5 ng/ml; NHE3: 1:1,000; VHA: 3 µg/ml in blocking buffer) at 4°C overnight. The next day, PVDF membranes were washed in TBS-T (3 x 10 min), incubated in blocking buffer with the appropriate secondary antibodies (1:10,000 in blocking buffer) for 1 h, and washed again in TBS-T (3 x 10 min). Bands were made visible through addition of enhanced chemiluminescence *Prime Western Blotting Detection Reagent* (GE Healthcare, Waukesha, WI) and imaged and quantified in a Universal III Hood (BioRad) using Image Lab software (version 6.0.1; BioRad). Following imaging, the PVDF membrane was incubated in Ponceau stain (10 min, RT) to estimate protein loading. Relative NKA, NHE3, and VHA protein abundance were normalized to the protein content within each lane.

### Immunohistochemistry

Rockfish gill samples were fixed in 4% paraformaldehyde in phosphate buffer saline (PBS, pH 7.4) on a shaker at 4°C overnight, immersed in 50% ethanol for 8 h, and stored in 70% ethanol at 4°C until processing. Fixed samples were rehydrated in PBS + 0.1% tween (PBS-T) for 5 min, rinsed in ice-cold PBS containing 1.5 mg/mL NaBH_4_ (6 x 10 min), then incubated in blocking buffer (PBS-T, 2% normal goat serum, 0.02% keyhole limpet hemocyanin) for 1 h at room temperature. Following this, samples were incubated with primary antibodies (NKA: 40 ng/mL; NHE3: 1:500; VHA: 6 µg/mL in blocking buffer) at 4°C overnight. On the next day, samples were washed in PBS-T, incubated with blocking buffer containing secondary antibodies (1:500) and the nuclear dye DAPI (1 µg/mL) for 1 h at room temperature. The stained gill samples were then washed in PBS-T, whole-mounted onto depressed glass slide fitted with a glass cover slip (No. 1.5, 0.17 mm), and imaged using a Zeiss inverted confocal microscope (AxioObserver Z1 with LSM800; Oberkochen, Germany) and Zeiss ZEN 2.6 blue edition software. To quantify gill NKA- and VHA-rich cell abundance, immunostained whole-mounted gill samples were imaged using 10x and 20x objective lenses (EC Plan-Neofluar 10x/0.30 Ph 1; Plan-Apochromat 20x/0.8 M27) and optimized settings for Alexa Fluor 488 (to visualize VHA; excitation 493 nm at 0.5% laser power, emission 517 nm, detection 510– 575 nm), Alexa Fluor 568 (to visualize NKA; excitation 577 nm at 0.2% laser power, emission 603 nm, detection 575–617 nm), and DAPI (to visualize nuclei; excitation 353 nm at 0.2% laser power, emission 465 nm, detection 410–470 nm). Z-stacks were produced as maximum intensity projection of ∼70–500 consecutive ∼1.93 µm optical sections. The average NKA- and VHA-rich ionocyte abundances were quantified in the proximal, medial, and distal sections of gill filaments (defined as in Christensen et al., 2012; Figure 1A) using FIJI *cell counter* plugin (Schindelin et al., 2012). From each of nine control and nine ERH-exposed rockfish (N=9), the numbers of NKA- and VHA-rich ionocytes were quantified after double-blind randomization in five lamellae from three filaments (total of 15 lamellae), which were averaged together for each replicate.

**Figure 1.**
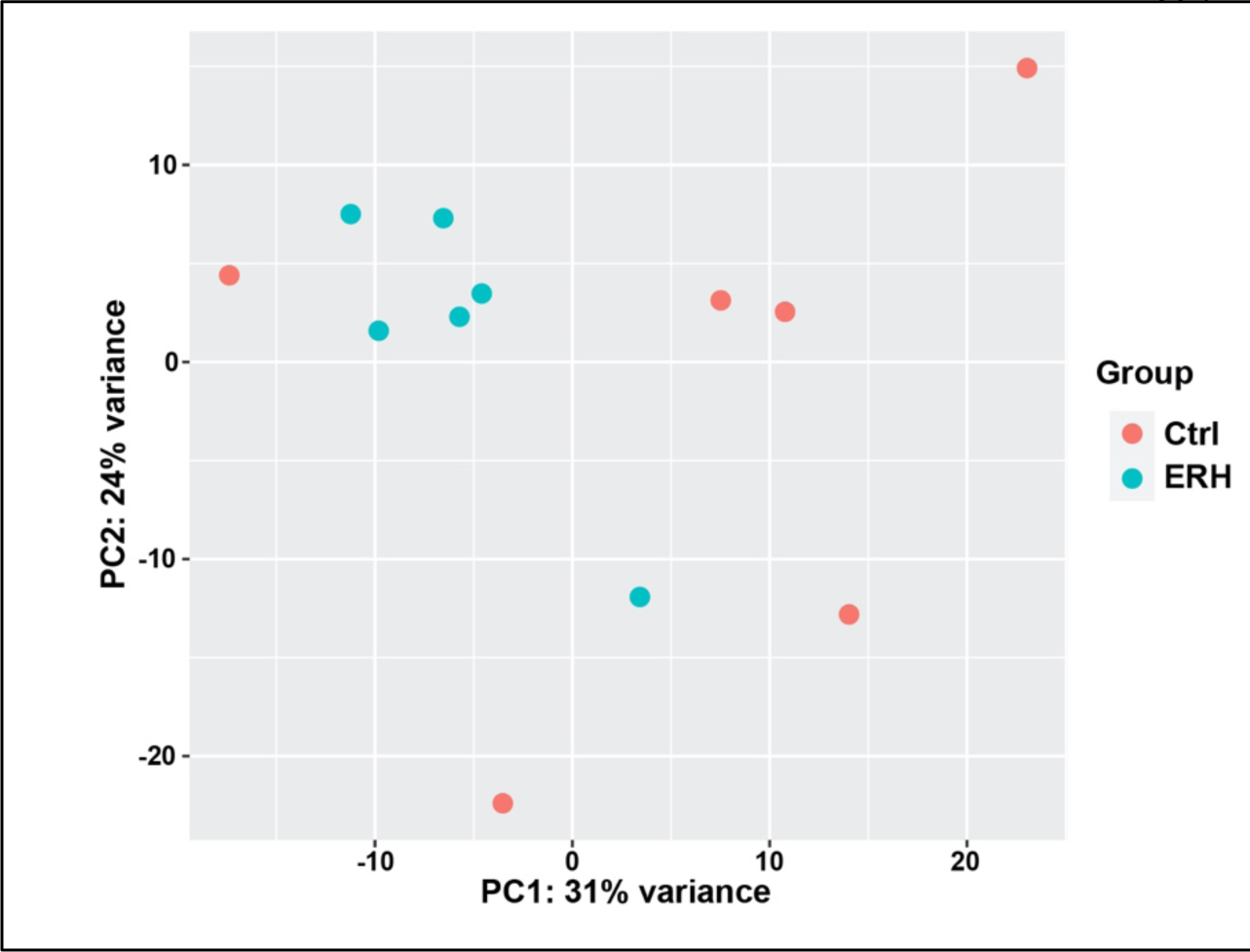
Principal component analysis of gene expression profiles. Gene expression visualized by the first two principal components (PC1: 31%, PC2: 24%). Each point represents an individual fish, and colors denote exposure to control (Ctrl; ∼570 µatm CO2, pH ∼7.9) or environmentally relevant hypercapnia (ERH; ∼1,600 µatm CO2, pH ∼7.5) conditions (N=6).

The subcellular localization of NKA, NHE3, and VHA was examined using 40x and 63x objective lenses (Zeiss LD LCI Plan-Apochromat 40x/1.2 lmm Korr DIC M27; Plan-Apochromat 63x/1.40 Oil DIC M27) and optimized settings for Alexa Fluor 488 (to visualize VHA or NHE3; excitation 493 nm at 0.6-1.0% laser power, emission 517 nm, detection 510–575 nm), Alexa Fluor 568 (to visualize NKA or NHE3; excitation 577 nm at 0.6-1.2% laser power, emission 603 nm, detection 575–617 nm), and DAPI (to visualize nuclei; excitation 353 nm at 0.6-1.0% laser power, emission 465 nm, detection 410–470 nm). Z-stacks of the gill ionocytes were visualized as maximum intensity projection of ∼70–250 consecutive ∼0.25 µm optical sections and analyzed using orthogonal cuts across the X-Z and Y-Z planes. Observations were conducted on five control and five ERH rockfish, on three filaments per rockfish.

### Scanning Electron Microscopy

Rockfish gill samples were fixed in 3% paraformaldehyde and 0.35% glutaraldehyde in 0.1 M cacodylate buffer overnight at 4°C and processed as described previously by Kwan *et al*. (2019). Briefly, the samples were dehydrated in 50% ethanol for 8 h and stored in 70% ethanol. On the day of processing, fixed gill samples were dehydrated in 100% tert-butyl alcohol applying 25% increments over 24 h, frozen in 100% tert-butyl alcohol at 4°C, and freeze dried using a VirTis benchtop freeze dryer (SP Industries, Gardiner, New York). Samples were then sputter-coated with gold and imaged using a ThermoFisher Apreo 1 with Apreo LoVac field emission gun under low-vacuum mode (20 kV, 1.6 nA, ∼0.03 mbar). Imaging was focused on the cells localized along the basal portion of the gill lamellae. The tasks of imaging, random assignment, and blind analysis of five ionocytes per rockfish (N = 8 control and N = 9 ERH) were performed by different investigators to ensure unbiased data collection.

### Statistical Analysis

Statistical tests were performed using R (version 4.0.3; R Development Core Team, 2013). Normalized RNAseq data were converted to the global Pearson’s inter-sample correlational coefficient and analyzed as distance values to infer relative differences between libraries, and to generate a hierarchical clustering profile using R and the *MDS* package. Principal component analysis (PCA) on the normalized data was conducted using DESeq2. Differentially expressed genes (DEGs; log2 fold-change ≥ 2) were identified based on significance (alpha of 0.05). Gene Ontology (GO) and Kyoto Encyclopedia of Genes and Genomes (KEGG) enrichment analysis was performed using gProfiler (biit.cs.ut.ee/gprofiler; Raudvere et al., 2019). Differences in variance of each gene were assessed using the Levene’s test.

Protein abundance, hematocrit, and apical morphology datasets were analyzed with the Student’s t-test (two-tailed). NKA- and VHA-rich ionocyte abundance were analyzed using two-way ANOVA with treatment and gill filament section as factors. Statistical assumptions of normality and homoscedasticity were assessed using the Shapiro-Wilk and F tests, respectively. Membrane-enriched NKA abundance required an inverse transformation, whereas membrane-enriched VHA abundance required a square root transformation to meet the parametric requirements. The SEM-based apical morphology dataset failed to meet the assumptions of normality even after transformation and was analyzed using the non-parametric Wilcoxon rank sum test (two-tailed) with continuity correction. An alpha of 0.05 was used for all datasets. Values are reported as means ± standard error of the mean (S.E.M).

## Results

We previously reported that rockfish exposed to ERH (∼1,600 μatm *p*CO_2_; pH ∼7.5) for three days completely regulated blood pH (ERH: 7.85 ± 0.04; Control: 7.75 ± 0.03) and that their blood plasma had significantly higher [HCO_3_^-^] (ERH: 5.16 ± 0.31 mM; Control: 2.37 ± 0.20 mM) (Kwan and Tresguerres, 2022). Here, we additionally report a significant ∼15% increase in haematocrit (Hct) in ERH-exposed rockfish (30 ± 1.0% vs. 26 ± 1.3% in controls).

Maintenance of blood pH combined with plasma HCO_3_ ^-^ accumulation indicated active and effective blood acid-base regulation in response to ERH. As a first step to elucidate the underlying mechanisms, we performed RNAseq in gills from control and ERH-exposed rockfish. The RNAseq data mapped to 24,577 genes annotated in the honeycomb rockfish genome and PCA revealed two components that together accounted for 55% of total variation in our dataset (PC1 = 31%, PC2 = 24%). The gene expression dispersal pattern in control rockfish was stochastic, whereas that in ERH-exposed rockfish was more tightly clustered together (Fig. 1).

However, only 94 genes (<0.4% of the total) were differentially expressed in ERH-exposed fish: 65 upregulated (Table S2) and 29 downregulated (Table S3). GO and KEGG analyses (false discovery rate ≤ 0.01) failed to identify differentially enriched pathways. Gene-by-gene analyses of DEGs revealed strong upregulation of genes coding for transcription factors and splicing regulators (RNA-binding motif 20, Zinc finger protein 239), for proteins implicated in angiogenesis and vascularization (neuropeptide Y receptor y2, acetylcholine receptor subunit α-7), and for others involved in cell adhesion and cytoskeletal remodeling (contactin 3a, collagen, laminin beta, calpain-5) (Table 2). However, the mRNA abundances of genes coding for proteins with established functions in fish acid-base and ion regulation were unchanged (Fig. 2A_1-3_; Fig. S2, S3), suggesting that blood acid-base regulation during ERH did not involve a transcriptional response. Alternatively, changes in mRNA levels could have taken place in the early phase of ERH exposure, affecting the abundance of acid-base regulatory proteins, and returned to baseline levels by the time of sampling on day 3. We investigated this possibility by quantifying the protein abundances of NKA, NHE3, and VHA_B_ in rockfish gills.

**Figure 2.**
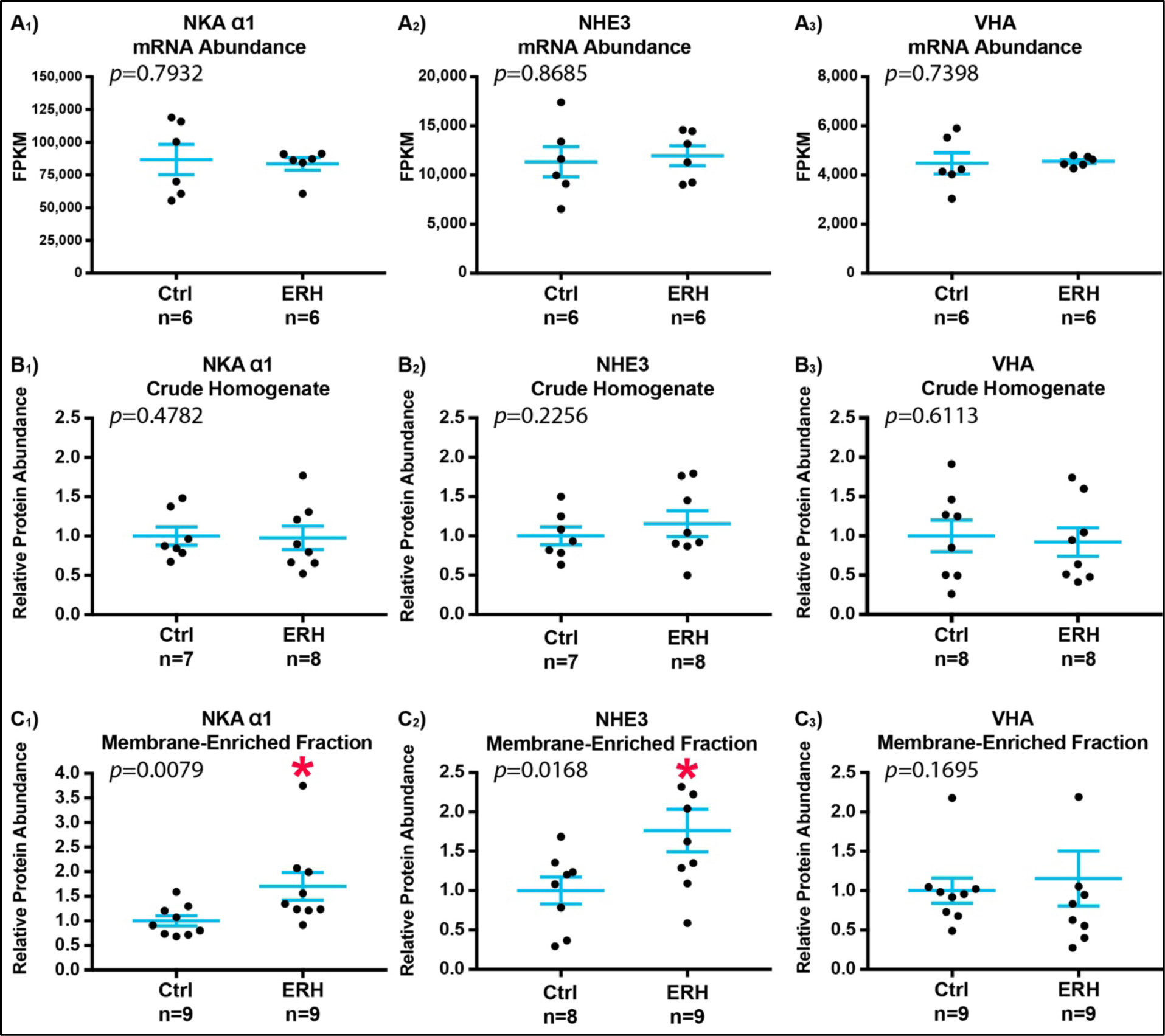
Gene expression and protein abundance in gills from control and ERH rockfish. Ctrl: fish exposed to control conditions (∼570 µatm CO2, pH ∼7.9). ERH: fish exposed to environmentally relevant hypercapnia (∼1,600 µatm CO2, pH ∼7.5). **A1-3)** mRNA abundance, **B1-3)** total protein abundance in crude homogenate (indicative of total abundance), and **C1-3)** protein abundance in the membrane-enriched fraction of **1**) Na^+^/K^+^-ATPase α subunit (NKA), **2**) Na^+^/H^+^ exchanger 3 (NHE3), and **3**) V-type H^+^ ATPase subunit B (VHAB). *Asterisk* denotes significant difference. Values are mean ± S.E.M. FPKM: Fragments Per Kilobase of transcript per Million mapped reads.

The total abundances of NKA, NHE3, or VHA_B_ protein in rockfish gill homogenates (estimated Western blots in gill crude homogenates) did not differ between control and ERH-exposed rockfish (Fig. 2B_1-3_), indicating a lack of translational response. In contrast, the abundances of NKA and NHE3 protein in gill cell membrane-enriched fractions were 50% higher in ERH-exposed rockfish than in control rockfish (Fig. 2C_1-2_), while VHA_B_ protein abundance in gill cell membrane-enriched fraction remained unchanged (Fig. 2C_3_). These results suggested that ERH-exposed rockfish translocated pre-existing NKA and NHE3 proteins into the plasma membrane of gill ionocytes to upregulate H^+^ excretion to seawater and regulate blood pH.

Next, we used immunohistochemistry to examine whether exposure to ERH affected the abundance of NKA-rich ionocytes and VHA-rich cells in rockfish gills. As previously reported for most other fish species (Evans et al., 2005), the NKA-rich ionocytes were much more abundant on the trailing edge than on the leading edge of rockfish gills (Fig. 3A-C). Strikingly however, the majority of these ionocytes were present along the bottom half of the gill lamellae (Fig. 3B, C), which contrasts with most other fishes studied so far, which typically have the ionocytes on the gill filament or at the base of the lamellae (Christensen et al., 2012; Skelton et al., 2024). The abundance of VHA-rich cells was approximately half of that of NKA-rich ionocytes, and they were present on both the leading and trailing edge of the filament as well as on the lamellae. A separate analysis along the proximal, medial, and distal portions of the gill filaments (see diagram in Figure 3A) revealed the NKA-rich ionocytes were significantly more abundant on the proximal and medial sections than in the distal section, while VHA-rich cells did not differ among the sections. These patterns were unaffected by ERH exposure (Fig. 3D, E and Table S4).

**Figure 3.**
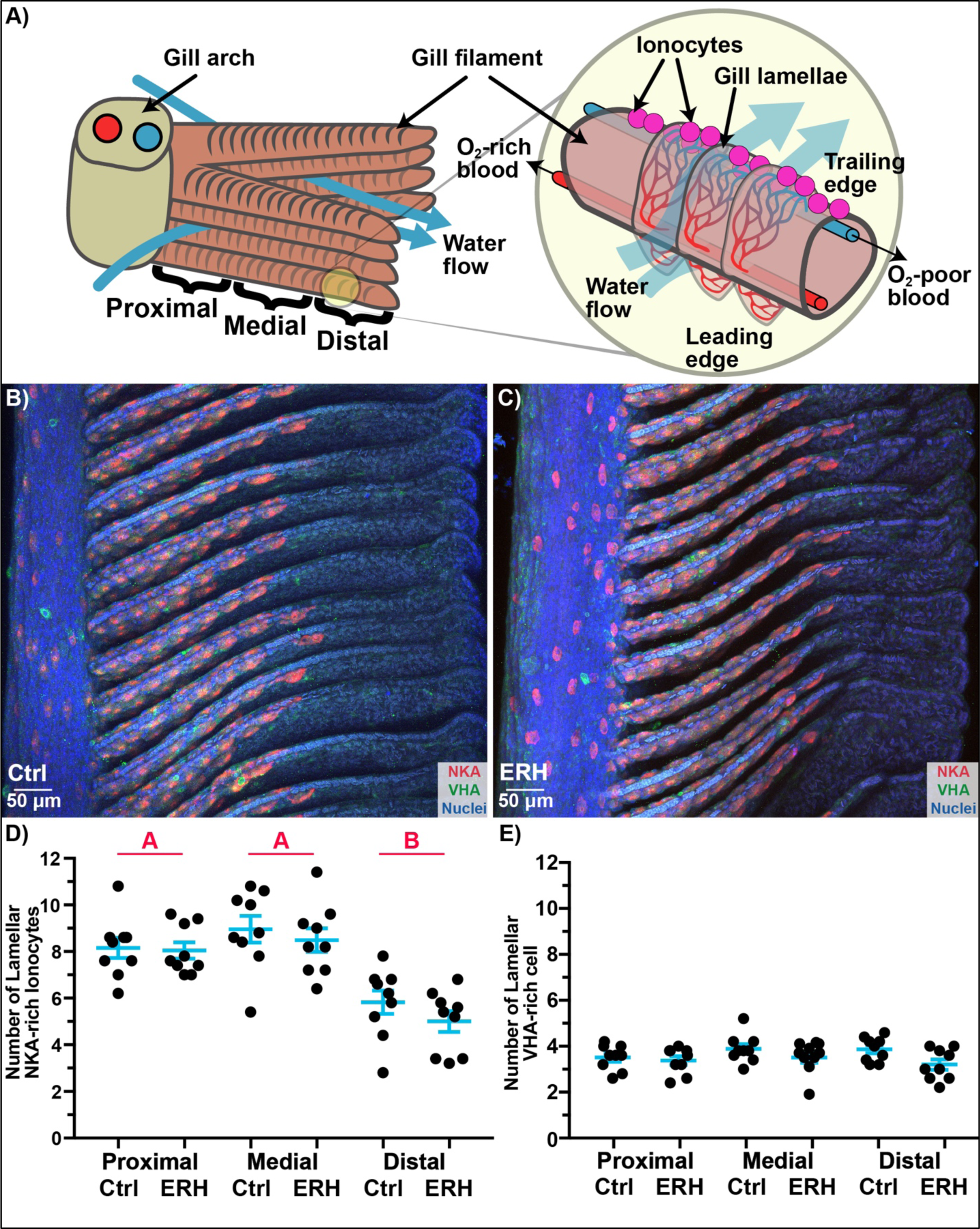
Na^+^/K^+^-ATPase-rich (NKA-rich) ionocytes and V-type H^+^-ATPase-rich (VHA-rich) cells in gills from control and ERH rockfish. **A)** Diagram of a fish gill arch, filaments, and lamellae showing ionocytes in the trailing edge (pink circles). **B, C)** Representative image of gills from control and ERH rockfish immunostained for NKA (red) and VHA (green). **D, E)** Abundance of NKA-rich ionocytes and VHA-rich cells along the lamellae at the proximal, medial, and distal sections of gill filaments. n=9. Values are mean ± S.E.M. Letters denote a significant difference in NKA-rich ionocyte abundance between filament sections; there were no significant differences between sections in VHA-rich cells, or between control and ERH treatments in either cell type.

Higher magnification 3D reconstructed confocal images of gill ionocytes revealed that NKA-rich ionocytes expressed NHE3 in the apical membrane (Fig. 4A, B), NKA adjacent to the apical membrane but in the basolateral membrane tubular system (Fig. 4), and VHA in the cytoplasm (Fig. 4C, D). Relative to control fish, ERH-exposed fish seemed to have a denser apical NHE3 signal and a NKA signal that extended towards the basal side of the ionocyte (Fig. 4B). These qualitative observations matched the Western blot results and, together, they indicate insertion of NHE3 into the apical membrane, and of NKA into the basolateral membrane of ionocytes. On the other hand, the VHA-rich cells had VHA along the basolateral edge of the cell, did not express NKA or NHE3, and these patterns were unaffected by the ERH exposure (Fig. S3).

**Figure 4.**
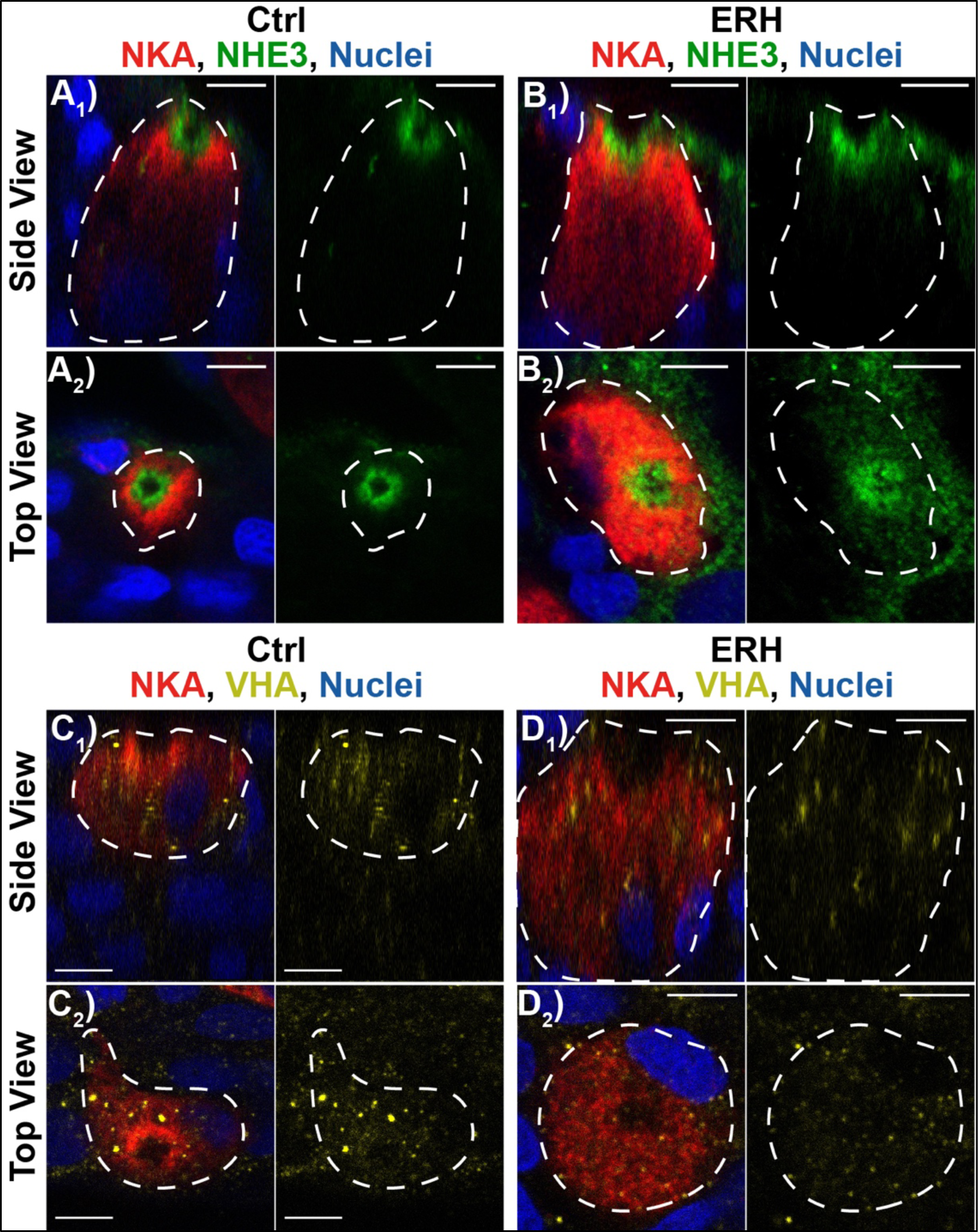
Protein localization in gill NKA-rich ionocytes from control and ERH rockfish. Na^+^/K^+^-ATPase (NKA), Na^+^/H^+^ exchanger 3 (NHE3), and V-type H^+^ ATPase (VHA) expression within NKA-rich gill ionocytes after exposure to A, B) control (ctrl) or C, D) environmentally relevant hypercapnia (ERH) conditions. A_1_-D_1_: confocal images along the side view (*XZ* plane); A_2_-D_2_: confocal images along the top view (*XY* plane). (n>5 per protein, per treatment). White dashed line indicates the boundary of the NKA-rich ionocyte, which were determined with transmitted photomultiplier (T-PMT). Scale bar = 5 μm.

Finally, we examined the apical morphology of rockfish gill ionocytes using SEM. In control fish, the majority of ionocytes had a smooth apical membrane surface (see example in Fig. 5A) and 15.0 ± 8.24% of the ionocytes that were randomly analyzed had extended apical microvilli (see example in Fig. 5B). Interestingly, ERH-exposed rockfish had 45.5 ± 10.54% of ionocytes with extended microvilli [or a significantly higher 3-fold increase (*p*=0.0253; Fig. 5C)].

**Figure 5.**
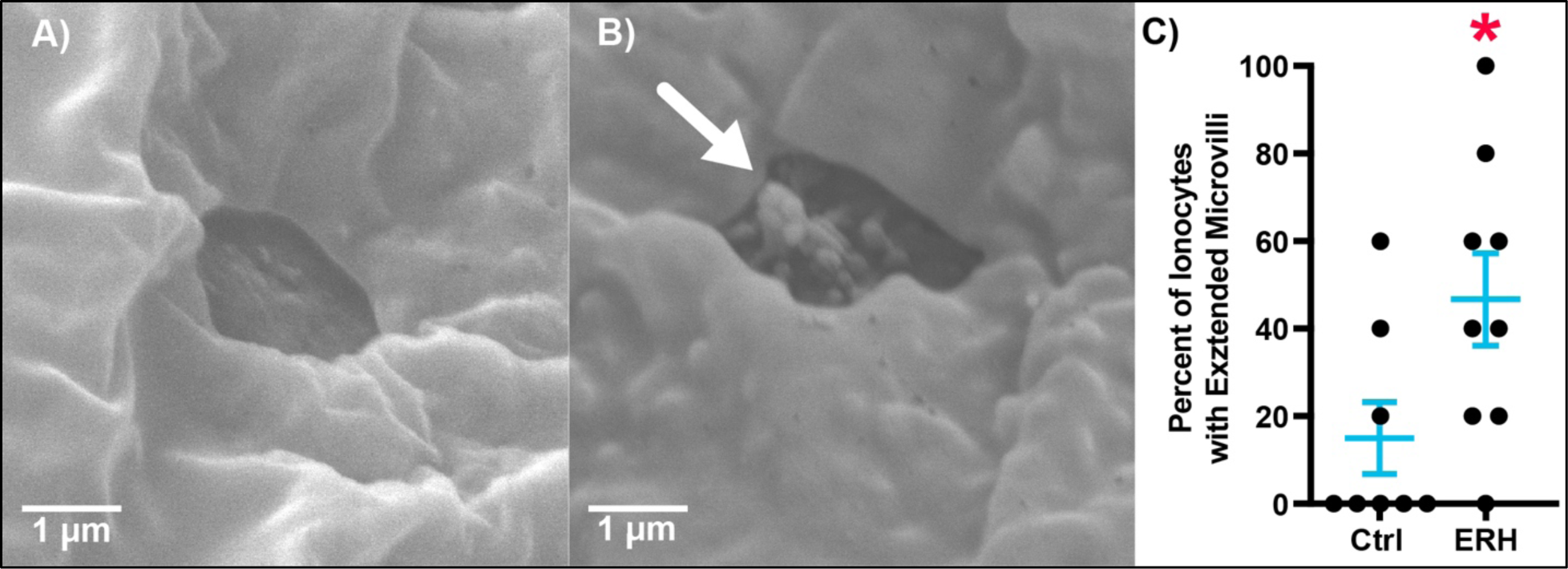
Apical morphology of gill ionocytes from control and ERH rockfish. The apical surface of gill ionocytes can have **A)** retracted or **B)** extended microvilli (white arrow). **C)** Blinded analysis of apical morphology revealed rockfish exposed to environmentally relevant hypercapnia (ERH) had significantly more ionocytes with extended microvilli than control (Ctrl) fish. *Asterisk* denotes significant difference. Ctrl n=8. ERH n=9. Values are mean ± S.E.M.

## Discussion

After three days of exposure to ERH at 1,600 μatm *p*CO_2_ (pH ∼7.5), rockfish blood pH was maintained at 7.85 and their plasma [HCO_3_^-^] more than doubled from ∼2.4 mM to ∼5.2 mM (Kwan and Tresguerres, 2022). These features are typical of active blood acid-base regulation upregulation in H^+^ excretion to seawater and in HCO_3_^-^ absorption into the blood following activation of gill ion-transporting proteins. *A priori*, various mechanisms could fill this role, including: (1) upregulation of mRNA abundance leading to increased protein abundance of ion-transporting proteins, (2) increased protein turnover (i.e. increased mRNA levels but constant protein levels), (3) increased protein abundance independent of mRNA changes (translational response), (4) posttranslational activation of preexisting proteins, or (5) combinations of these mechanisms. Additionally, the upregulation of H^+^ excretion and HCO_3_^-^ could have been related to a proliferation of gill ionocytes, which could confound the interpretation of the mRNA and protein quantification experiments. A systematic exploration ruled out all these options but one: the insertion of preexisting NKA and NHE3 into the basolateral and apical membranes of NKA-rich ionocytes coupled with microvilli extension to expand apical surface area (Fig. 6). Additional posttranslational modifications on NKA, NHE3 and other ion-transporting proteins, such as phosphorylation (Tresguerres et al., 2019), are likely and deserve further examination.

**Figure 6.**
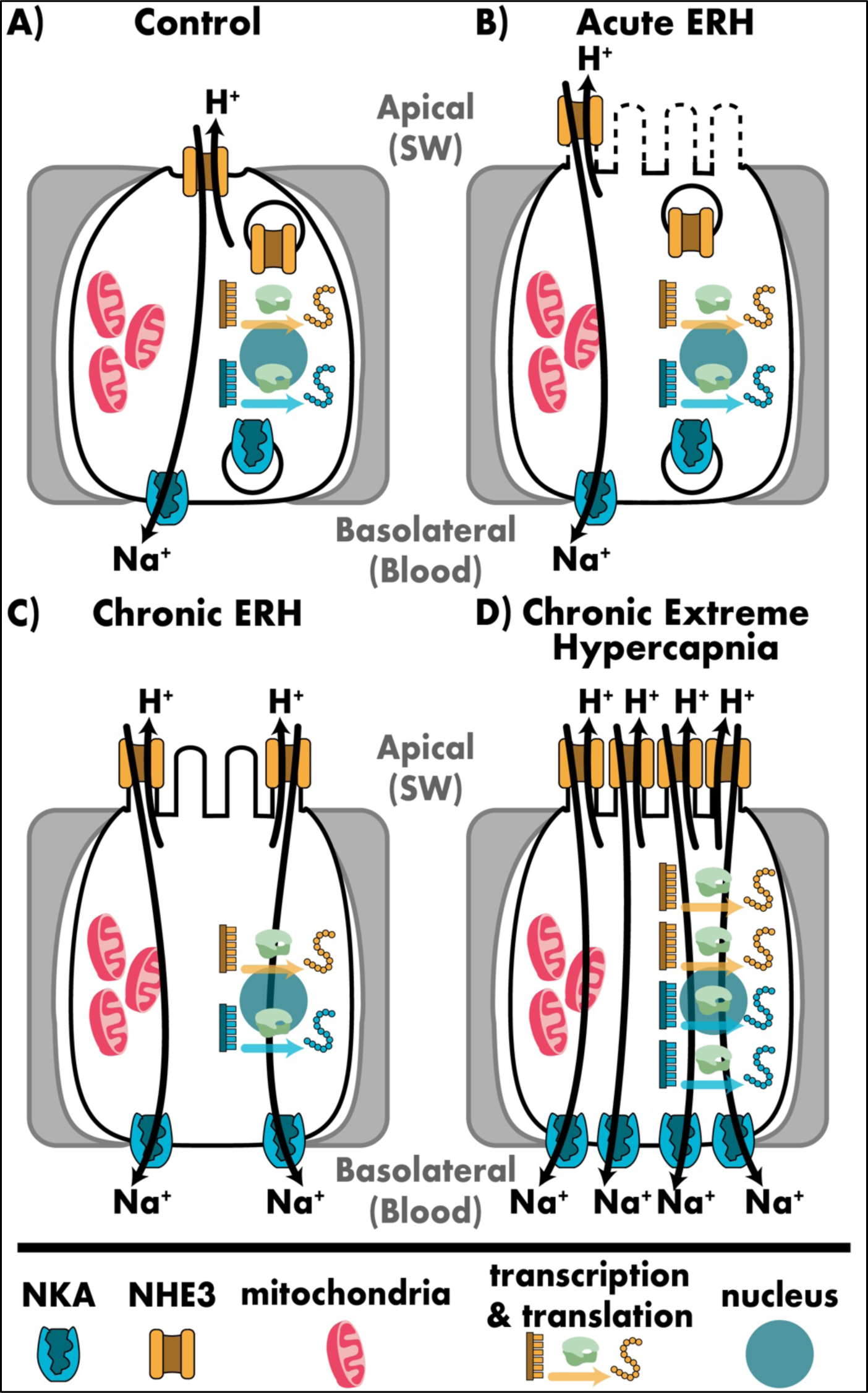
Putative mechanisms by which marine fish gill ionocytes regulate blood acid-base in response to hypercapnic challenges of different magnitude and duration. A) Under control conditions, the ionocytes have a baseline rate of H^+^ excretion driven by basolateral Na^+^/K^+^-ATPase (NKA) and apical Na^+^/H^+^-exchanger 3 (NHE3); there are reserve pools of both transporters in the cytoplasm (potentially in vesicles) and gene transcription and translation are at baseline levels. **B)** During an acute environmentally relevant hypercapnia (ERH), H^+^ excretion is upregulated *via* a widening of the ionocyte apical membrane. Other posttranslational regulatory mechanisms such as microvilli extension (dotted lines) and phosphorylation are likely. **C)** During a chronic ERH, H^+^ excretion is further upregulated *via* insertion of the reserve pools of NKA and NHE3 into the basolateral and apical membranes, respectively, and by apical microvilli extension. **D)** During a chronic extreme hypercapnia, H^+^ excretion is upregulated even further by increased transcription and translation of NKA and NHE3 (and possibly also other acid-base relevant proteins such as carbonic anhydrases and Na+/HCO3-cotransporters). The additional NKA and NHE3 readily insert in the ionocyte plasma membrane.

The lack of significant changes in VHA mRNA, total protein, and membrane protein abundance, VHA-rich cell abundance, and VHA translocation to the apical membrane of any gill cell type all but rule out a role of gill VHA in excreting H^+^ to seawater to compensate an ERH-induced blood acidosis. The roles of VHA in the gills of marine teleosts remain unknown. In the NKA-rich ionocytes, is it is unclear whether VHA is present in vesicles, in the basolateral membrane tubular system, or both. While it is possible that VHA absorbs H^+^ into the blood to compensate a postprandial blood alkalosis as shown in elasmobranchs (Roa et al., 2014; Tresguerres et al., 2005), it is not known if rockfish even experience a postprandial alkalosis. Alternatively, the observed VHA localization might correspond to lysosomes and play a role in digesting biological polymers for energy metabolism and cell structural maintenance. In the VHA-rich cells, VHA is localized at the basolateral edges; however, the goblet shape of these cells and their ubiquitous presence throughout the gill epithelium resemble the mucus cells described in fish skin (Ottesen and Olafsen, 1997) and gills (Alkan and Oğuz, 2021; Mir and Channa, 2011) and that may suggest a role in mucus production rather than in ion transport.

In addition to the gill responses, we found that ERH-exposed rockfish had a significant ∼15% increase in Hct compared to controls, which resembles a previous report for European sea bass after acute exposure to pronounced hypercapnia (10,000 μatm *p*CO_2_, pH ∼6.90 for 2h) (Montgomery et al., 2022). Potential sources of this increased Hct are unknown but include erythropoiesis, adrenergic spleen contraction that releases red blood cells into the systemic circulation. Likewise, the effect of this increased Hct on blood gas transport capacity, acid-base regulation, and cardiac output remain to be determined. The gill mRNA expression profile remained remarkably stable in the ERH-exposed rockfish, as only 94 out of 24,577 genes (<0.4%) were differentially expressed and no GO or KEGG pathways were differentially enriched. Noteworthy, rockfish exposed to ERH for 72h exhibited no significant changes in mRNA abundance of any genes coding for acid-base regulatory proteins. However, the distinct PCA clustering of gill samples from ERH-exposed rockfish indicates a more homogenous general transcriptomic profile compared to that of controls, which suggests a physiological response to the ERH. The upregulation of many genes involved in angiogenesis and vascularization hints to links with the significantly higher Hct of ERH-exposed rockfish, and perhaps also with their increased plasma [HCO_3_^-^]. However, these highly speculative possibilities must be followed up with protein quantification and functional experiments to establish the potential significance of gene upregulation on rockfish physiology and their responses to ERH.

A synthesis of studies of marine fishes reveals a multi-tier gill ionocyte response to regulate blood pH in response to hypercapnia of increasing magnitude and duration (Fig. 6). The core acid-excreting mechanism in the gill ionocytes of marine fishes is based on basolateral NKA and apical NHE3. A pool of each of these transporters is constitutively membrane-located and actively excreting H^+^ that is produced by routine metabolism, while other pools are ‘on reserve’ (Fig. 6a). The acute response to hypercapnia involves an increase of the NHE3-containing apical surface area (Montgomery et al 2022) (Fig. 6b). This response is likely to be involved in the compensation of a blood acidosis resulting from burst swimming thus explaining why highly active fishes tend to be able to regulate blood pH in response to hypercapnia at a faster rate than more sluggish species (Melzner et al., 2009b; Montgomery et al., 2022; Tresguerres et al., 2020). The chronic acid-base regulatory response depends on the magnitude of the hypercapnia.

During an ERH, fish additionally insert the reserve pools of NKA and NHE3 into the ionocyte basolateral and apical membranes, respectively, and extend its apical microvilli to increase the surface area for NHE3-dependent H^+^ excretion (Fig. 4, 5, 6c). Like the acute response, the chronic response to ERH does not require the upregulation of ion-transporting proteins (Fig. 2, Allmon and Esbaugh, 2017; Esbaugh et al., 2012; Michael et al., 2016; Shrivastava et al., 2019). In contrast, the chronic response to severe hypercapnia (>6,000 μatm *p*CO_2_; pH <7.00) additionally requires increased transcription and translation of NKA and NHE3 (Deigweiher et al., 2008; Edwards et al., 2005; Melzner et al., 2009a) (Fig. 6d). In all these cases, upregulation of H^+^ excretion *via* the NKA-NHE3 machinery in gill ionocytes allow fishes to fully regulate their blood pH and compensate for the initial acidosis. However, H^+^ excretion to seawater is accompanied by proportional absorption of HCO_3_^-^ into plasma, and HCO_3_^-^ accumulation in fish plasma has been linked to otolith overgrowth (Kwan and Tresguerres, 2022), increased skeletal biomineralization (Di Santo, 2019), neurobehavioral disturbances (Tresguerres and Hamilton 2017), and protection of cardiac function (Lo et al., 2020). Additionally, HCO_3_^-^ plays many important roles in mammalian physiology (Cordat and Casey, 2009), which further highlights the need to study the effects of increased [HCO_3_^-^] resulting from compensation of hypercapnia-induced blood acidosis on fishes. Another important consideration is that while ERH at nighttime in highly populated areas and during upwelling is associated with hypoxia, hypercapnia due to anthropogenic OA is not. This highlights the need for future experiments examining the interactions between environmental O_2_ and CO_2_/pH levels on the acid-base regulatory and metabolic responses of aquatic animals.

## Conclusion

The use of RNAseq, Western blotting on crude homogenates and membrane-enriched fractions, immunolocalization, SEM, and measurements of blood acid-base parameters allowed us to identify that rockfish exposed to an ERH regulate their blood pH by recruiting pre-existing NKA and NHE3 into the basolateral and apical membranes of gill ionocytes, respectively. In contrast, no significant changes in gene or protein expression were involved. This integrative physiological genomic approach exemplifies the need to explore physiological responses to ERH (and more broadly, to changing environments) at multiple levels of biological organization spanning from gene expression, protein abundance and intracellular localization, whole whole-animal physiology. Additionally, our study highlights that homeostatic responses are tuned to the magnitude of the environmental or metabolic changes, and that such tuning can be tracked down to the cellular level. These considerations are particularly important when studying the effects of OA on fishes, as the underlying acid-base disturbance is relatively minor, and the associated compensatory responses may not be readily detectable using standard techniques.

## Acknowledgement

This research was supported by National Science Foundation (NSF) grant IOS #1754994 to M.T, as well as by NSF Graduate Research Fellowship and NSF Postdoctoral Research Fellowship in Biology (award #1907334) to GTK. We thank Phil Zerofski (SIO) for collecting the fish and helping with aquarium matters. We are grateful to Taylor Smith, Shane Finnerty, and Gabriel Lopez for their help with aquarium maintenance and fish care. The authors would also like to thank Dr. Junya Hiroi (St. Marianna University School of Medicine, Kawasaki, Japan) for generously donating the anti-NHE3 antibodies, and Sabine Faulhaber (UC San Diego) for technical assistance with the scanning electron microscope.

