## Supplemental Material for "Gill ionocyte remodeling mediates blood pH regulation in rockfish (*Sebastes diploproa*) exposed to environmentally relevant hypercapnia"

**Table S1:** General RNAseq analysis metrics in splitnose rockfish (*Sebastes diploproa*) pre- and post- quality control.

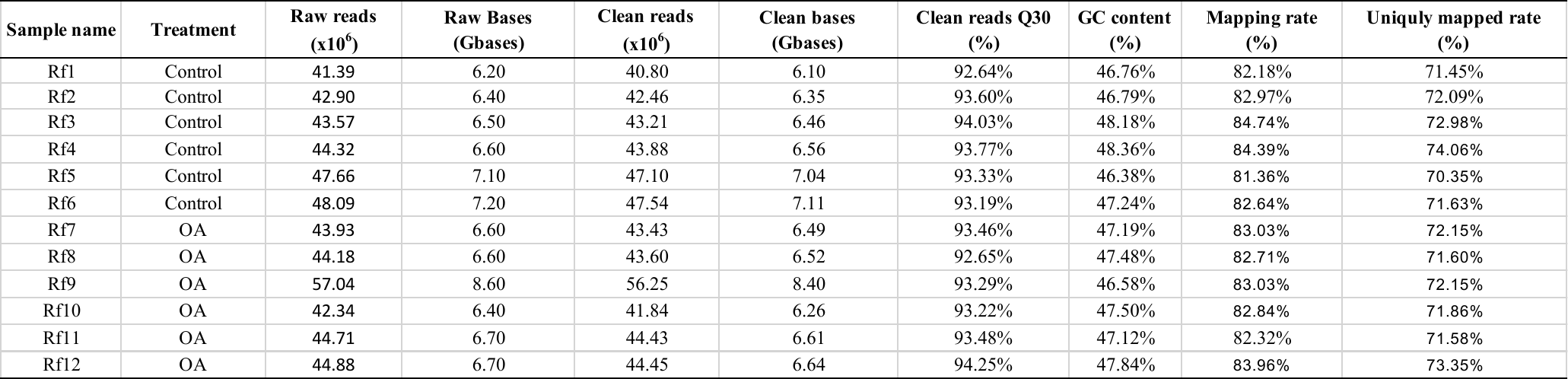

**Table S2:** Differentially expressed genes in gills from rockfish that were upregulated following ERH exposure.

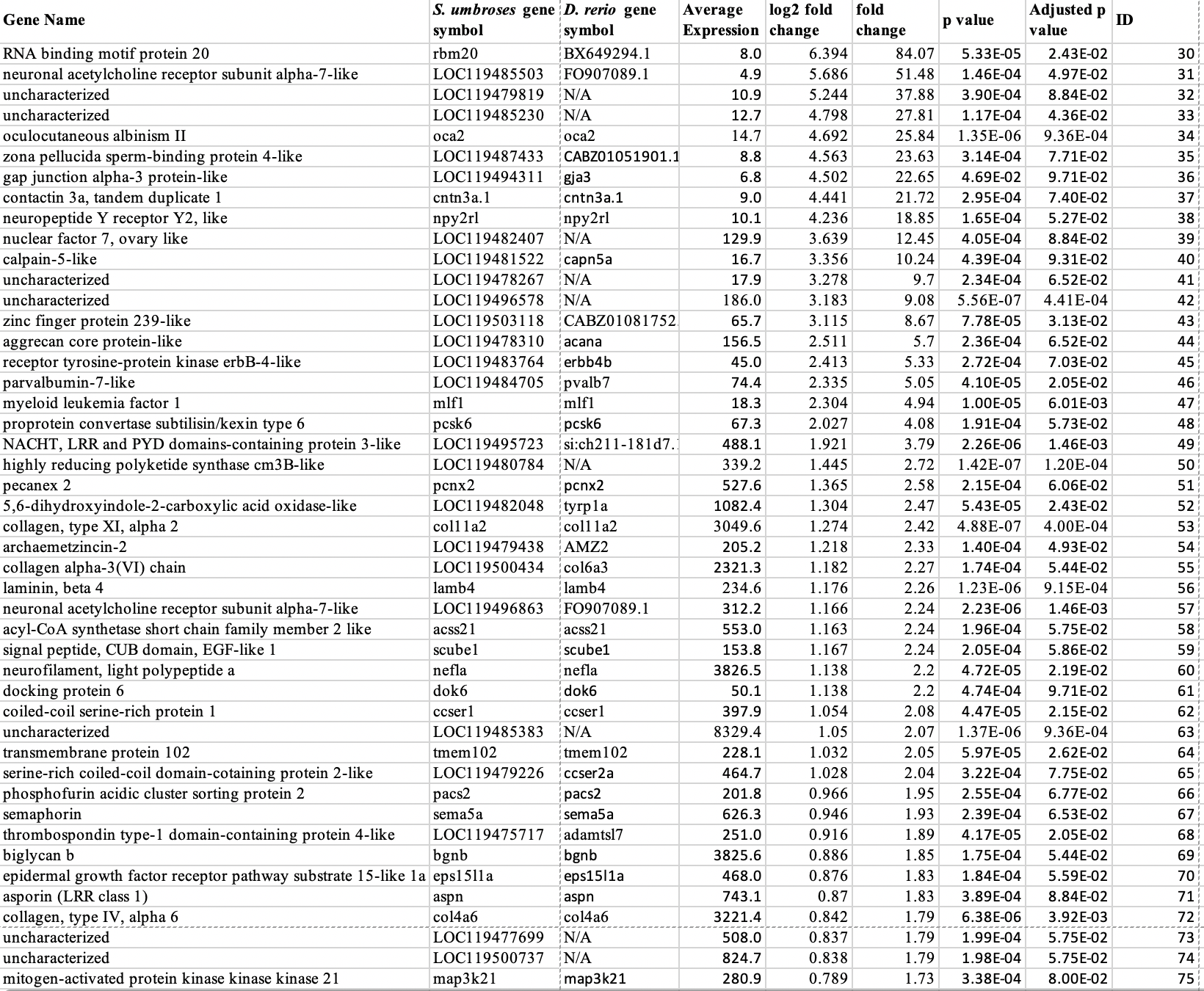

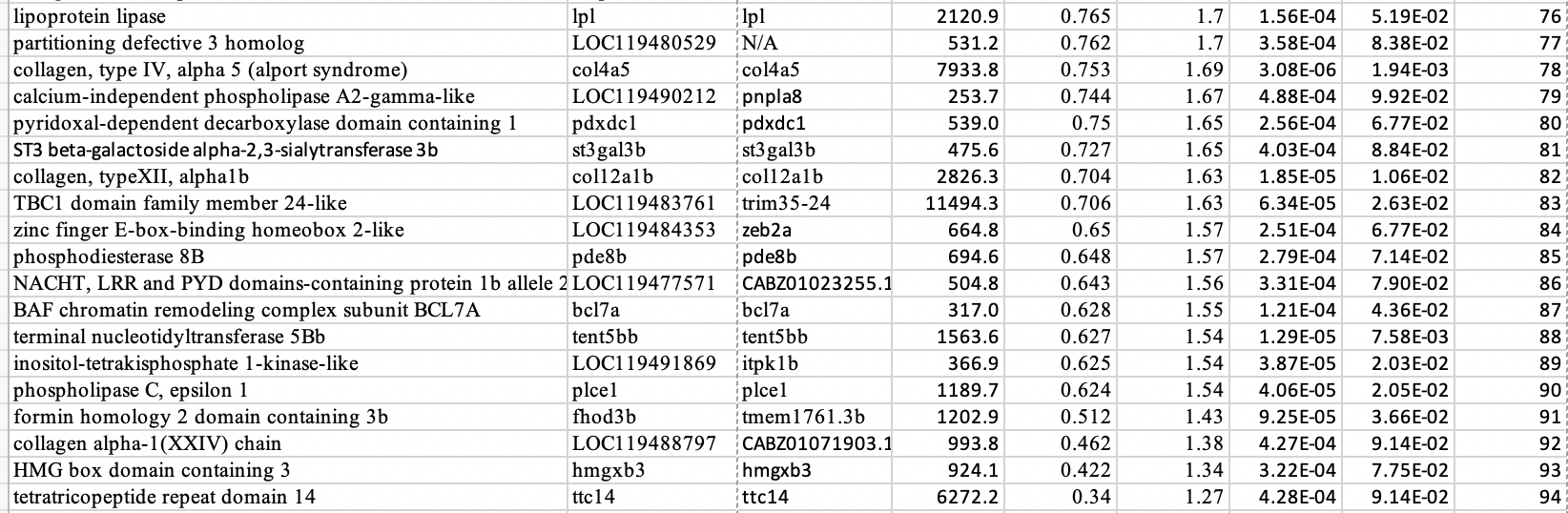

**Table S3:** Differentially expressed genes in gills from rockfish that were downregulated following ERH exposure.

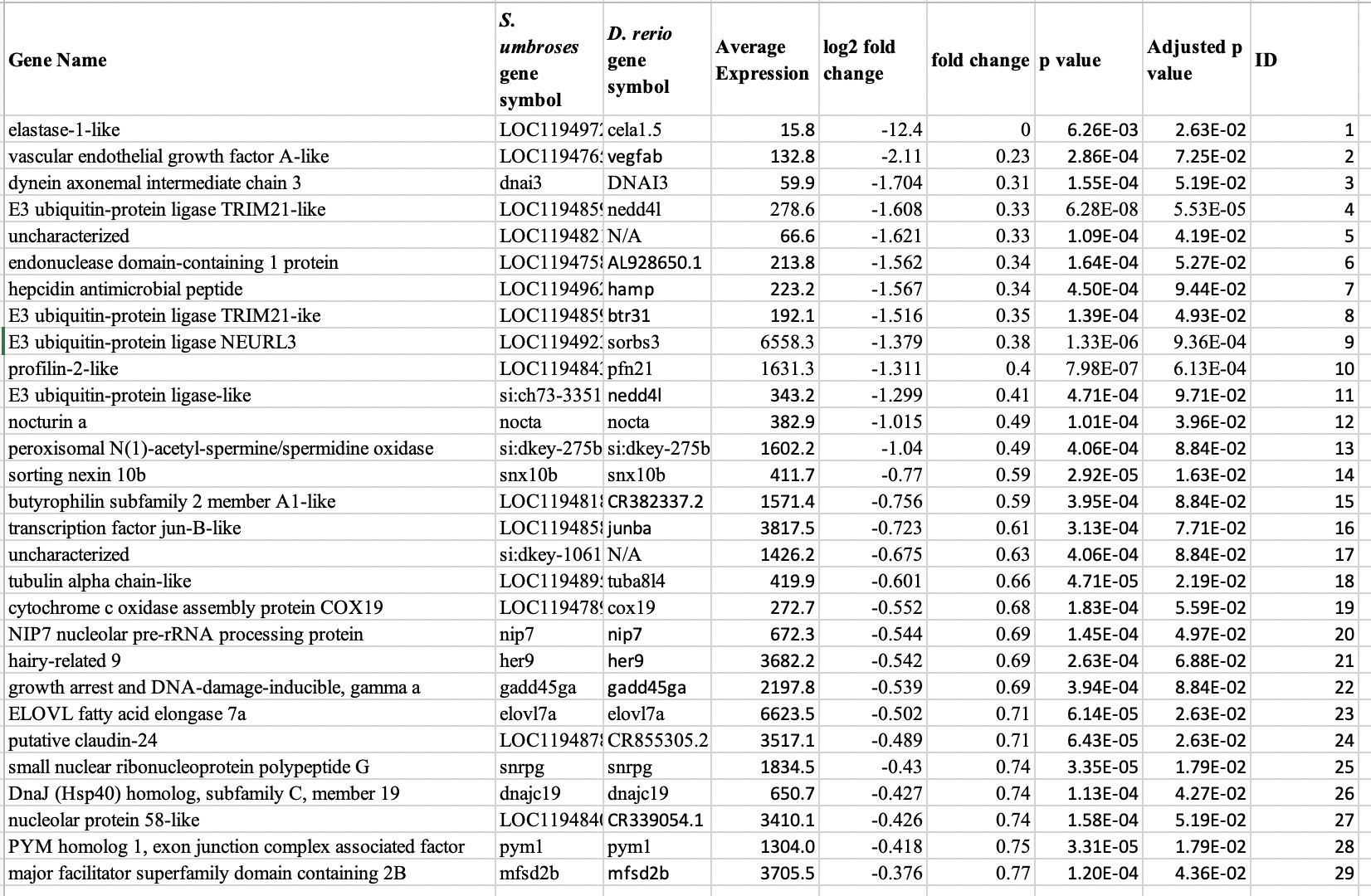

**Table S4: Number of Na^+^/K^+^-ATPase-rich (NKA-rich) ionocytes and v-type H^+^-ATPase-rich (VHA-rich) cells along the lamellae in different sections of gill filaments from control and ERH rockfish**. Values are mean ± S.E.M. N=9. Letters denote a significant difference in NKA-rich ionocyte abundance between filament sections; there were no significant differences between VHA-rich cells or between control and ERH treatments for either cell type. The data in this Table corresponds to Figure 3.

|  | Control | | | ERH | | |
| --- | --- | --- | --- | --- | --- | --- |
| Filament section | Proximal | Medial | Distal | Proximal | Medial | Distal |
| NKA-rich Ionocyte | 8.2±0.4^A^ | 9.0±0.6^A^ | 5.8±0.5^B^ | 8.0±0.4^A^ | 8.5±0.5 ^A^ | 5.0±0.4^B^ |
| VHA-rich Cell | 3.5±0.2 | 3.9±0.2 | 3.9±0.2 | 3.4±0.2 | 3.8±0.2 | 3.2±0.2 |

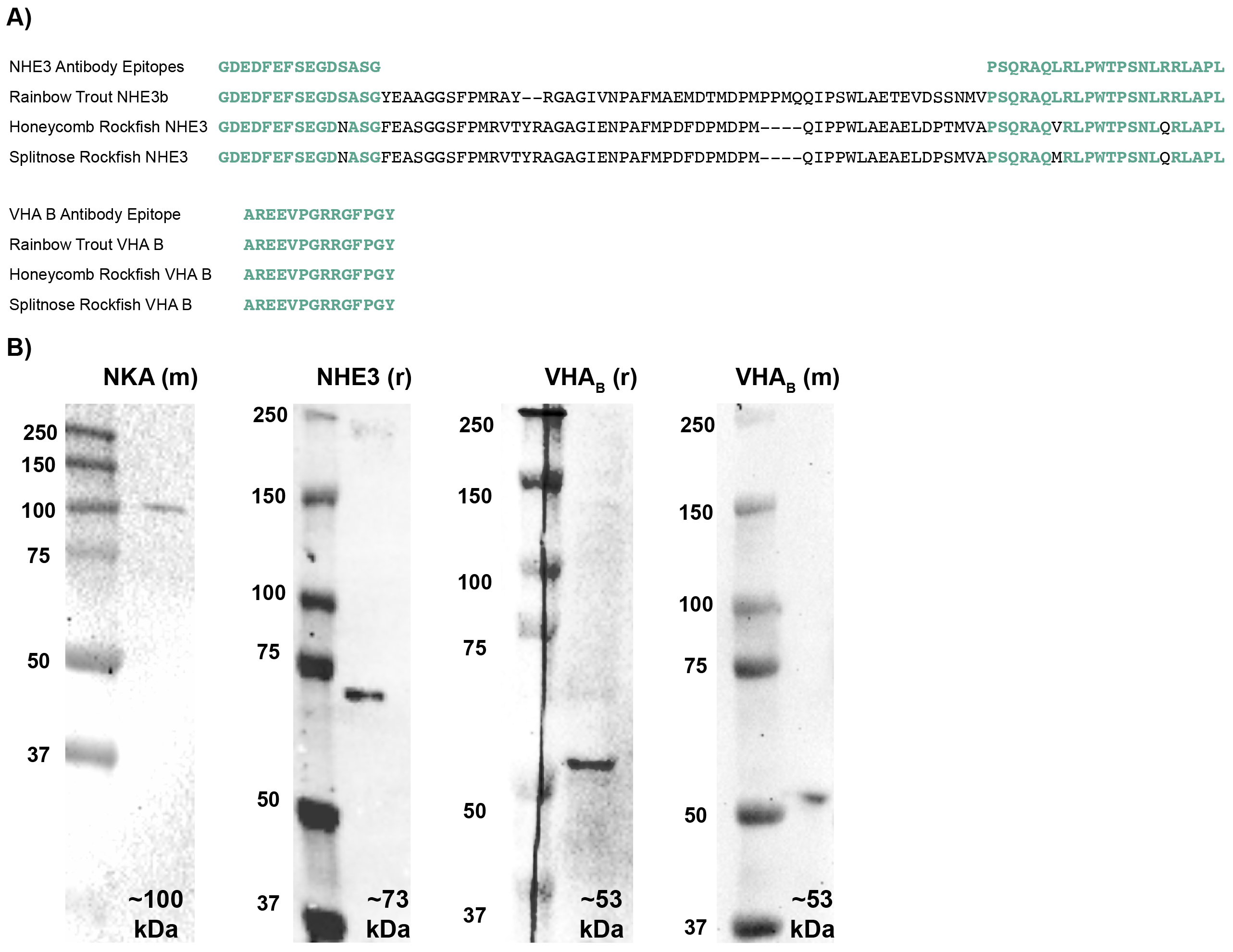

**Figure S1: Antibody validation. A)** Epitope regions (in green) recognized by the anti-NHE3 and anti-VHA_B_ antibodies; notice the high conservation in splitnose rockfish. **B)** Western blotting on gill homogenates detected distinct bands matching the predicted size of each protein. NKA (m): mouse monoclonal antibody against Na^+^-K^+^-ATPase subunit α; NHE3 (r): rabbit polyclonal antibodies against rainbow trout NHE3; VHA_B_ (r): rabbit polyclonal antibodies against VHA subunit B; VHA (m): mouse monoclonal antibody against VHA_B_. See Methods for more details about the antibodies.

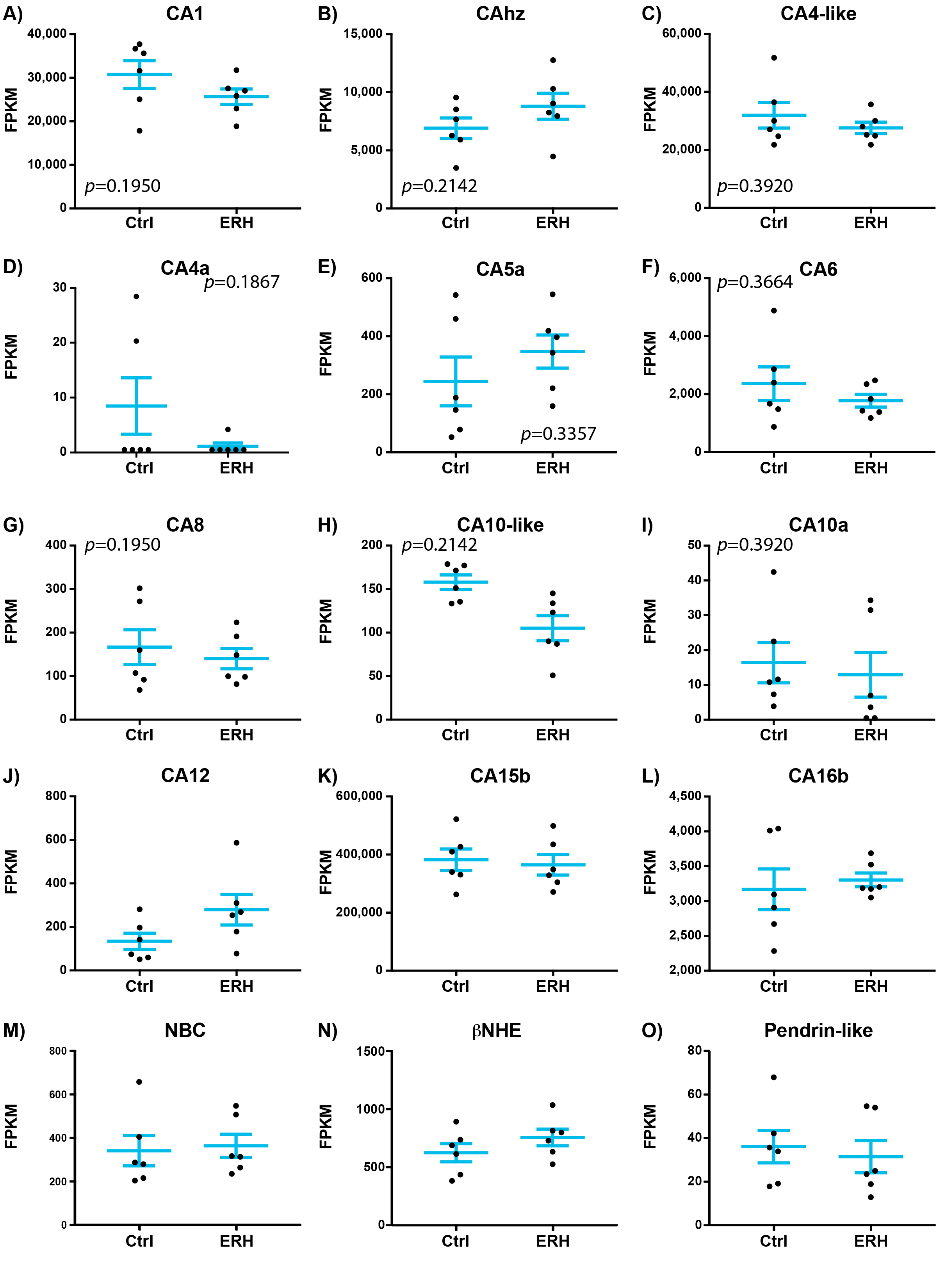

**Figure S2:** **Expression of acid-base relevant genes in gills from control and ERH rockfish.** FPKM: Fragments Per Kilobase of transcript per Million mapped reads. CA: carbonic anhydrase, NBC: Na^+^/HCO_3_^-^ co-transporter (slc4a4a), βNHE: beta-Na^+^/H^+^ exchanger (slc9a1b), Pendrin: Cl^-^/HCO_3_^-^ exchanger (slc26a4). Values are mean ± S.E.M. N=6.

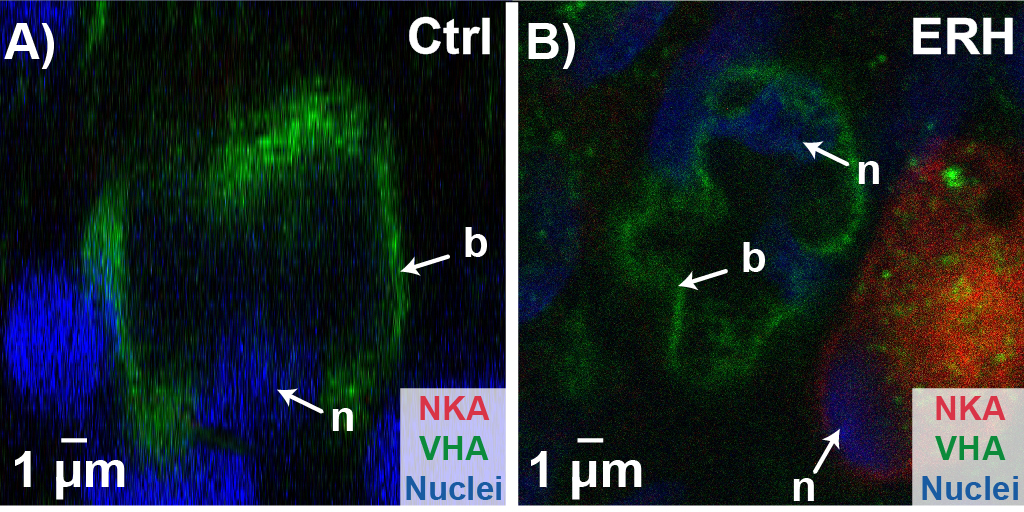

**Supplemental Figure 3**: **Protein localization in gill VHA-rich cells from control and ERH rockfish.** V-type H^+^ ATPase (VHA) expression within VHA-rich cells after exposure to A) control (ctrl) or B) environmentally relevant hypercapnia (ERH) conditions. NKA: Na^+^/K^+^-ATPase.
